# Bundle sheath cell-dependent chloroplast movement in mesophyll cells of C_4_ plants analyzed using live leaf-section imaging

**DOI:** 10.1101/2024.09.13.612856

**Authors:** Yuta Kato, Takao Oi, Yoshikatsu Sato, Mitsutaka Taniguchi

**Affiliations:** Graduate School of Bioagricultural Sciences, Nagoya University, Nagoya 464-8601, Japan; Institute of Transformative Bio-Molecules, Nagoya University, Nagoya, 464-8601 Japan

**Keywords:** Aggregative movement, bundle sheath cell, C_4_ photosynthesis, *Eleusine coracana*, mesophyll cell, chloroplast movement, live imaging

## Abstract

In C_4_ plants, mesophyll (M) chloroplasts aggregate toward bundle sheath (BS) cells in response to environmental stress, which would contribute to C_4_ photosynthetic cycle adjustment between M and BS cells. However, it remains unclear whether mesophyll chloroplast movement is an intercellular response mediated by BS cells. One major challenge to resolving this is the difficulty in observing how M chloroplasts aggregate toward adjacent BS cells due to scattering and absorption of observation light in live-leaf tissues. We established a live leaf-section imaging technique that enables the long-term observation of sections of chemically unfixed leaf blades, with which we quantitatively analyzed M chloroplast movements. Another challenge in clarifying the contribution of BS cells to M chloroplast movement is the selective ablation of BS cells without impairing their function of M cells. To investigate the necessity of BS cells for M chloroplast movement, we developed a method to remove BS cells only based on differences in shape and size between M and BS cells. We also found that chloroplasts in M cells without adjacent BS cell contents did not show typical aggregative movement but showed a light-avoidance response. This indicates that the M chloroplast aggregative movement occurs during communication with BS cells.

**Highlight:** We established live leaf-section imaging to observe individual chloroplast movements in multi-layered cells and found that bundle sheath cells are involved in the aggregative movement of mesophyll chloroplasts.

## Introduction

C_4_ plants have leaves composed of multiple cell layers, including two types of photosynthetic cells, mesophyll (M) and bundle sheath (BS) cells. These two cell types form a concentric arrangement called “Kranz anatomy” (Brown, 1975). The C_4_ photosynthetic cycle is driven across both cell types and serves as a pump to concentrate and supply CO_2_ to Rubisco within the BS chloroplasts (Hatch, 1999), thereby improving photosynthetic efficiency by suppressing photorespiration (Sage *et al*., 2012). In C_4_ leaves, M chloroplasts are fewer than in C_3_ leaves and lack Rubisco, whereas BS chloroplasts are larger than M chloroplasts and contain Rubisco (Stata *et al*., 2014). Regarding the intracellular arrangement of chloroplasts, chloroplasts in the BS cells are located in either a centripetal (close to the vascular tissue) or centrifugal (close to the M cells) position, depending on the plant species (Dengler and Nelson, 1999). In contrast, M chloroplasts are discretely distributed along the plasma membrane throughout the cell under no-stress conditions. In contrast, they aggregate toward the BS side of M cells in response to environmental stresses, such as drought, salinity, hyperosmosis, and intense light, whereas BS chloroplasts hardly move under these stress conditions (Lal and Edward, 1996; Yamada *et al*., 2009; Maai *et al*., 2011b; Taniguchi and Cousins, 2018; Maai *et al*., 2020; Kato *et al*., 2022; 2023). The directional movement of C_4_ M chloroplasts is known as “aggregative movement”. This chloroplast movement depends on actin filaments and induced by blue light but not red light (Kobayashi *et al*., 2009; Yamada *et al*., 2009; Maai *et al*., 2011b). Chloroplast aggregative movement is thought to contribute to the efficiency of intercellular metabolite transport and the remobilization of leaked CO_2_ by shortening the distance between M chloroplasts and BS cells under stress conditions (Yamada *et al*., 2009; Maai *et al*., 2011a; 2020; Kato *et al*., 2023). However, there is no evidence suggesting that the movement of chloroplasts in M cells is due to intercellular communication with BS cells.

Methods for studying chloroplast movement in live plants are well established, particularly for analyzing chloroplast photo-relocation movements in response to light intensity and wavelength (Senn, 1908; Wada *et al*., 2003; Wada, 2013; Wada and Kong, 2018). Transmission measurements and band assays (Inoue and Shibata, 1973; Kagawa *et al*., 2001; Wada, 2013), polarized light irradiation (Haupt and Scheuerlein, 1990), and microbeam irradiation (Wada *et al*., 1983; Wada, 2013) have been used to elucidate the mechanisms and significance of chloroplast movement. However, these techniques are challenging to apply to chloroplast aggregative movements, which include moving in the vertical direction relative to the leaf surface within the multilayered, complex structures of C_4_ leaves. Therefore, previous studies on chloroplast aggregative movements in C_4_ plants have relied on observing leaf cross-sections prepared after chemical fixation (Yamada *et al*., 2009; Kato *et al*., 2023). This method has not accurately captured chloroplast movement in the living state. There has long been a need to develop a technique to observe how M chloroplasts move toward BS cells within complex leaf tissue to elucidate the mechanism of chloroplast aggregative movement.

In this study, we developed a useful live leaf-section imaging method that analyzes individual behaviors of M chloroplasts in C_4_ plants. Using this method, we quantitatively analyzed M chloroplasts based on their distance from adjacent BS cells. Furthermore, we succeeded in preparing sections that excluded BS cells using a precise sectioning technique and investigated M chloroplast behavior in the absence of BS cells. Based on these investigations, we found that the aggregative movement of M chloroplasts requires intercellular communication with BS cells.

## Materials and Methods

### Plant material and growth conditions

Seeds of finger millet (*Eleusine coracana* L. Gaertn.) was purchased from Snow Brand Seed Co. Ltd., Japan, sterilized in 5% (w/v) sodium hypochlorite solution for 5 min, and imbibed on wet filter paper. Seeds were germinated in a growth chamber under a 14 h photoperiod (08:00–22:00) with a temperature of 28/20 ℃ (day/night). The PPFD was approximately 400 µmol m^-2^ s^-1^ with metal halide lamps (MF400GC/BUD, Koito Co. Ltd., Japan). The relative humidity was maintained at more than 60% all day. Germinated seedlings were then transplanted into 100 mL pots filled with culture soil (Hanachan Baiyodo Premium; Hanagokoro, Japan) based on bark compost mixed with coconut fiber, humic minerals, and Kanuma soil and grown in the same chamber for approximately two weeks. The uppermost fully expanded leaves (seventh leaf) were used in the experiments, and all treatments were started between 09:30 and 10:30.

### Live leaf-section imaging

The 5 mm square segments were excised from the leaf blades and vacuum-infiltrated for 5 min with purified water. The leaf segments were embedded in 5% (w/v) melted agar. The agar was chilled and stirred during the embedding process until immediately before solidification to prevent heat damage to the leaf segments. The leaf segment embedded in the agar was cut into a 100 µm thick transverse section using a vibrating blade microtome (VT1200S; Leica Microsystems, Germany). The obtained leaf section was carefully moved in a water droplet made with the surface tension of water at the tips of tweezers and placed in a narrow groove filled with 0.1% agar solution in clear silicone rubber on a glass slide (Supplementary Figure S1A). The adaxial or abaxial side of the leaf section was made to face the incident direction of blue light (Supplementary Figure S1B). Agar was added to the mounting solution to reduce fluidity and prevent the shifting of leaf sections during observation. In addition to the narrow groove in which the leaf section was placed, a large square groove was incorporated into the silicone rubber (Supplementary Figure S1A). This large square groove reduces the flow of the solution and prevents the leaf section from flowing out of the groove when a cover glass is used. The mounted leaf section was observed at 25°C under red observation light (< 1 µmol m^-2^ s^-1^) and snapped per 1 min with a microscope (BX51, Olympus, Japan) equipped with a CMOS camera (DP74; Olympus, Japan) (Supplementary Figure S1C). Red light filters that cut light below 600 nm were placed after the halogen lamp and in front of the camera in the optical path of the microscope light. Blue LED light (500 µmol m^-2^ s^-1^, maximum peak at 450 nm; ISL-150X150-BB45, CCS Inc., Japan) irradiated to the side of the mounted sample. As the position of the leaf section easily changed immediately after setting the cover glass, microscopic pre-observations were conducted for at least 1 h to ensure that the shift of the leaf section stabilized before starting the experiments.

### Removal of contents of BS cells and observation of chloroplast movement

Small segments of leaf blade embedded in agar were cut into longitudinal sections with intact M cells at 40 µm thickness to remove the BS cell contents or 60 µm thickness to keep the BS cell contents using the vibrating blade microtome. From these longitudinal sections, those containing vascular tissue were selected for subsequent live imaging, as described above.

### Quantification of M chloroplast arrangement

We calculated the relative distance (*RD*) of the M chloroplasts from the BS side to evaluate the degree of chloroplast aggregation at each time point, as previously described (Kato *et al*., 2023; Supplementary Figure S4A). In this study, we used the *RD* value as an index of the position of each chloroplast in an M cell, as calculated from the digital images of the sections. The pixel coordinates of the points were measured using Katikati Counter software (GTSOFT, Japan).

### Quantification of M chloroplast moving

To perform a tracking analysis of the movement of each chloroplast, we utilized the “Manual Tracking” plugin available in Fiji (Schindelin *et al*., 2012). Tracking intervals were every minute, and tracking was aborted when each chloroplast was no longer recognizable because of overlap with other chloroplasts or was out-of-focus because of chloroplast movement. In addition, we obtained “*Total movement*” to quantify the extent of chloroplast movement in any direction and “*Displacement*” to quantify the extent of chloroplast movement toward the BS cells during 1 h after blue light irradiation (Fig. 3A). In addition, *Displacement/Total movement* was calculated to represent the proximity to the BS cell side. Correlations were determined between these parameters and chloroplast position at the initial of blue light irradiation “*Initial position*” to determine if the distance from BS cells affects the movement of M chloroplasts. The numbers of measured chloroplasts and other information are presented in the figure legends.

### Statistical analyses

Data were analyzed using Microsoft Excel with the add-in software Statcel 3 (OMS Publishing, Japan). The significance of differences among the medians of *RD* values for each treatment was estimated using Student’s *t*-test or the Tukey–Kramer method. The importance of differences among M chloroplast motility for each treatment was estimated using the Mann–Whitney *U* or Steel–Dwass tests. The correlation between M chloroplast motility and chloroplast position at the initial treatment stage was estimated using Pearson’s correlation coefficient. The details of the statistical parameters are provided in the figure legends.

## Results

### Establishment of live leaf-section imaging to continuously observe chloroplast movement in the C_4_ plants, Eleusine coracana *L. Gaertn*

To observe chloroplast movement within mesophyll tissues in C_4_ plants, a transverse section of the *E. coracana* leaf blade, without chemical fixation, was set in a narrow groove in a transparent silicone rubber on a glass slide and irradiated with blue light from the adaxial side to induce chloroplast movement (Supplementary Figure S1). This process has three critical points: handling of sliced leaves, immobilization of leaves during preparation, and the light irradiation method. First, preserving the leaf morphology and avoiding damage from drying or contact was crucial when leaf sections were moved from the microtome stage to their proper position in the grooves on glass slides. Therefore, the leaf sections were transferred using water droplets formed by the surface tension of water at the tips of tweezers. It is also essential to prevent sample drift during prolonged observations. To address this issue, we modified the grooves’ shape and increased the mounting solution’s viscosity by adding agar (Supplementary Figure S1A, B). Transverse sections of the living leaf tissue maintained a typical Kranz anatomy; M cells surrounded the BS cells with a ring-like layer (Fig. 1). M chloroplasts were dispersed throughout the cell, whereas the BS chloroplasts occupied centripetal positions on the vein side of the BS cell without light stimulation (Fig. 1A). Finally, the responses to the observation and stimulus lights should be distinguished. Therefore, we used red light, which does not induce chloroplast movement, for observation, whereas blue light was used to induce chloroplast movement from the epidermal side of the leaves using an external light source (Supplementary Figure S1C). Time-lapse observations demonstrated that the M chloroplasts aggregated toward adjacent BS cells upon blue light irradiation (Supplementary Movie S1), whereas they did not show this aggregative movement in the absence of blue light (Supplementary Figure S2A; Supplementary Movie S2). Quantitative analysis of the time-lapse images showed that the aggregative response reached a maximum at approximately 4 h and was maintained even after 16 h (Supplementary Figures S3A, S4). In contrast, BS chloroplasts maintained their positions centripetal to the vein throughout the experimental period. When the blue irradiation was stopped after 4 h, the aggregated chloroplasts gradually dispersed to a state similar to that before blue irradiation (Supplementary Figure S5A; Supplementary Movie S3). These results indicated that our leaf-section imaging method unequivocally minimized the effects of slicing injury and enabled, for the first time, sufficiently long observations of chloroplast movement in C_4_ plants.

**Fig. 1.**
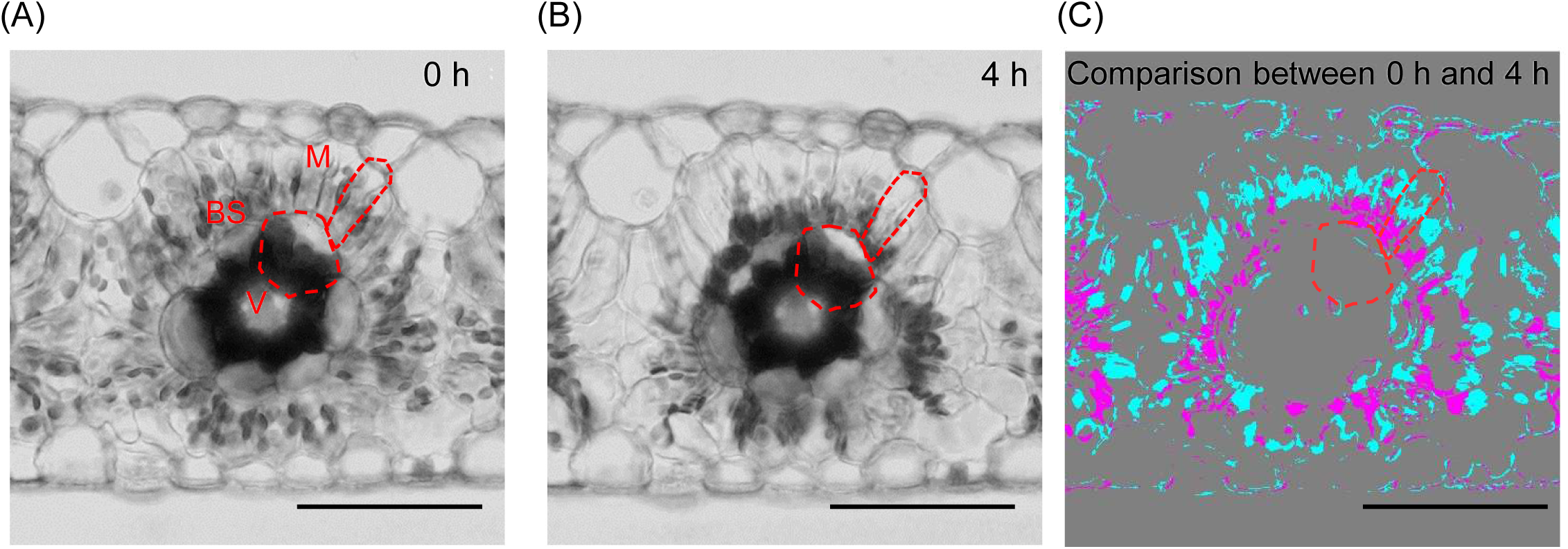
Continually observation changes in M chloroplast arrangement within the same leaf tissue in response to blue light. Images of transverse leaf tissue from finger millet leaf blades. The leaf section was irradiated with blue light (500 µmol m^-2^ s^-1^) from the adaxial side. The upper part of each image shows the adaxial side. (A, B) Images were obtained after blue light irradiation for 0 and 4 h. (C) Comparison between 0 and 4 h after blue light irradiation. The images in (A) and (B) were binarized, and the areas that appeared and were lost after 4 h are shown in magenta and cyan, respectively. Areas that did not change are shown in gray. This image was created using the “Make Binary” and “Image Calculator – Subtract” functions in ImageJ. M, mesophyll cell; BS, bundle sheath cell; V, vascular bundle. Bars = 50 µm.

### Similar responsiveness of chloroplast movement in the adaxial and abaxial leaf tissues

Focusing on the light incident direction on the leaf, M chloroplasts on the adaxial side of the leaf tissues moved remarkably closer to the BS cells under blue light irradiation from the adaxial side (Supplementary Movie S1). In contrast, those on the abaxial side moved remarkably closer to the BS cells when blue light irradiation was applied from the abaxial side (Supplementary Movie S4). These results indicate no significant difference in the reactivity of M cells between the adaxial and abaxial sides of the leaf. Instead, the direction of light incidence strongly influences them, and M cells on the side of light incidence show high responsiveness.

### Blue light accelerates the speed of M chloroplast movement toward BS cells

To quantitatively evaluate how M chloroplasts move toward the BS cells, we tracked individual M chloroplasts from time-lapse images acquired every minute (Fig. 2; Supplementary Movie S5; Supplementary Figures S2B, C; S3B; S5B, C). We quantitatively evaluated the chloroplast movements in each experiment for 1 h when the chloroplasts moved actively. In the blue light irradiation experiment (Supplementary Movie S1), 1 h after the initial irradiation; in the no blue light irradiation experiment (Supplementary Movie S2), 1 h after the initial observation; and in the turn-off of blue light irradiation experiment (Supplementary Movie S3), 1 h after the irradiation was turned off. We found that the speed of M chloroplast movement toward BS cells was significantly faster than that in the absence of blue light irradiation and turn-off of blue light irradiation (Table 1). Although this trend was more pronounced on the adaxial side, it was similar on both sides. Therefore, subsequent experiments and analyses were conducted only on the adaxial sides of the leaf sections.

**Fig. 2.**
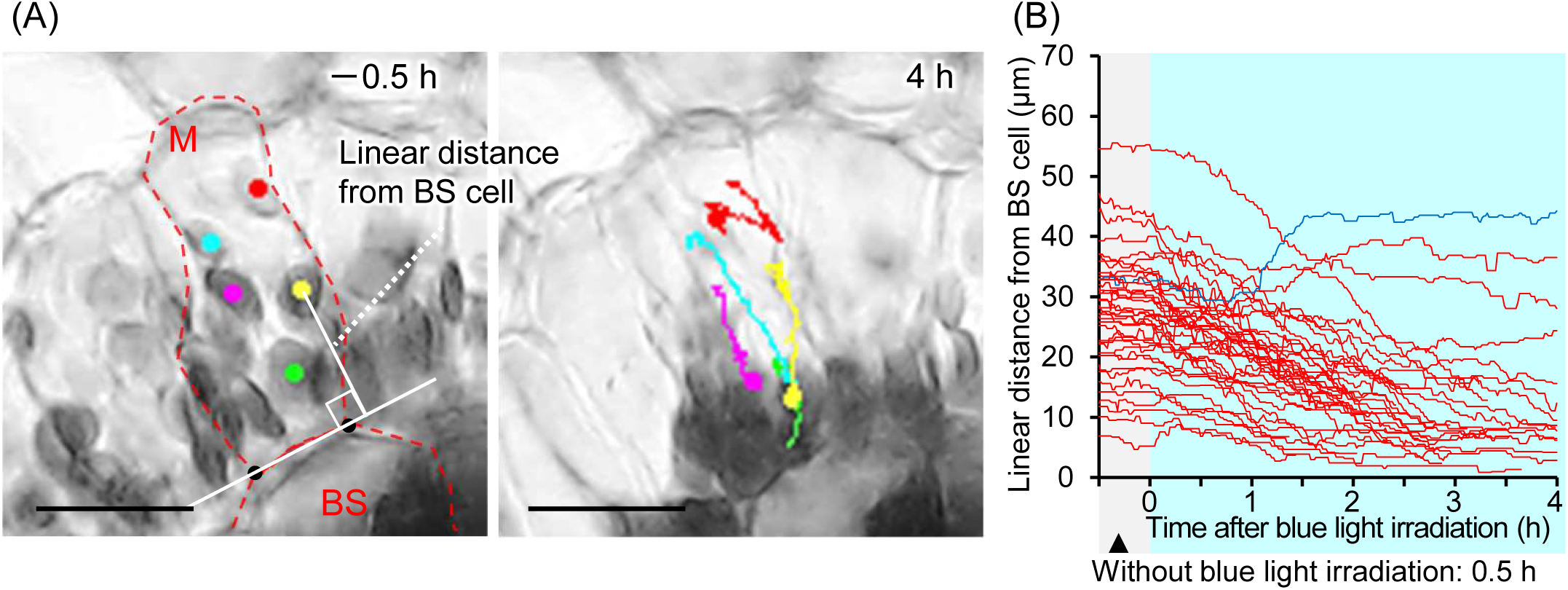
Temporal change in the distance of each M chloroplast from BS cell with blue light irradiation. The leaf section was irradiated with blue light (500 µmol m^-2^ s^-1^) from the adaxial side of the leaf section. (A) Images of adaxial M cells at the initial observation (−0.5 h) and 4 h after irradiation. A dot indicates the chloroplast position at each observation time, and chloroplast movement up to that time is indicated by a line. (B) Temporal changes in linear distance from BS cells in each chloroplast at each time were plotted every minute from 0.5 h before to 4 h after the blue light irradiation. If the chloroplasts could not be tracked because they overlapped or became out of focus, tracking was terminated at that point. Red lines represent chloroplasts that moved closer to the BS cells, and blue lines represent those that moved farther from the BS cells after 4 h of blue light irradiation compared to their initial position at the initial stage of irradiation (t = 0). The line colors were determined based on the tracking endpoint for chloroplasts that could not be tracked until 4 h of blue light irradiation. More than ten chloroplasts were randomly selected for monitoring from the three individuals’ adaxial side of the leaf sections (chloroplast number = 39). M: mesophyll cells; BS: bundle sheath cells. Bars in (A) = 25 µm.

**Table 1.** M chloroplast motility for 1h.

|  | Blue light | No irradiation | Turn-off<br>after 4 h blue light |
| --- | --- | --- | --- |
| **Adaxial side** |  |  |  |
| Speed of chloroplast movement ($\mu\text{m min}^{-1}$) | 0.28±0.14 a | 0.04±0.02 c | 0.09±0.09 b |
| *Displacement/Total movement* | 0.42±0.24 a | 0.00±0.54 c | -0.52±0.66 b |
| **Abaxial side** |  |  |  |
| Speed of chloroplast movement ($\mu\text{m min}^{-1}$) | 0.20±0.16 a | 0.04±0.02 c | 0.07±0.04 b |
| *Displacement/Total movement* | 0.23±0.32 a | -0.31±0.45 b | -0.21±0.49 b |
| **Both sides** |  |  |  |
| Speed of chloroplast movement ($\mu\text{m min}^{-1}$) | 0.24±0.15 a | 0.04±0.02 b | 0.08±0.14 b |
| *Displacement/Total movement* | 0.35±0.28 a | -0.16±0.52 b | -0.38±0.60 b |

Each value represents the mean ± SD. Each value was calculated 1 h after the initial irradiation in the blue light irradiation experiment, 1 h after the initial observation in the no blue light irradiation experiment, and 1 h after irradiation was turned off in the turn-off of blue light irradiation experiment. Different letters indicate a significant difference at $p < 0.05$ (Steel–Dwass test, adaxial side chloroplast numbers = 35-39. abaxial side chloroplast numbers = 33-36).

In addition, we investigated whether the aggregation of M chloroplasts was influenced by their initial distance from adjacent BS cells. The correlation between chloroplast behavior and distance from the BS cells at the initial irradiation (*Initial position*) was analyzed. It was found that there were weak positive correlations between *Initial position* and *Total movement* or *Displacement*, but not *Displacement/Total movement* (Fig. 3). These trends indicate that M chloroplasts that were located farther from the BS cells and near the light source appeared to move more toward the BS cells. The distance from the BS cells did not seem to affect the rate of approaching toward the BS cells in terms of total migration.

**Figure 3.**
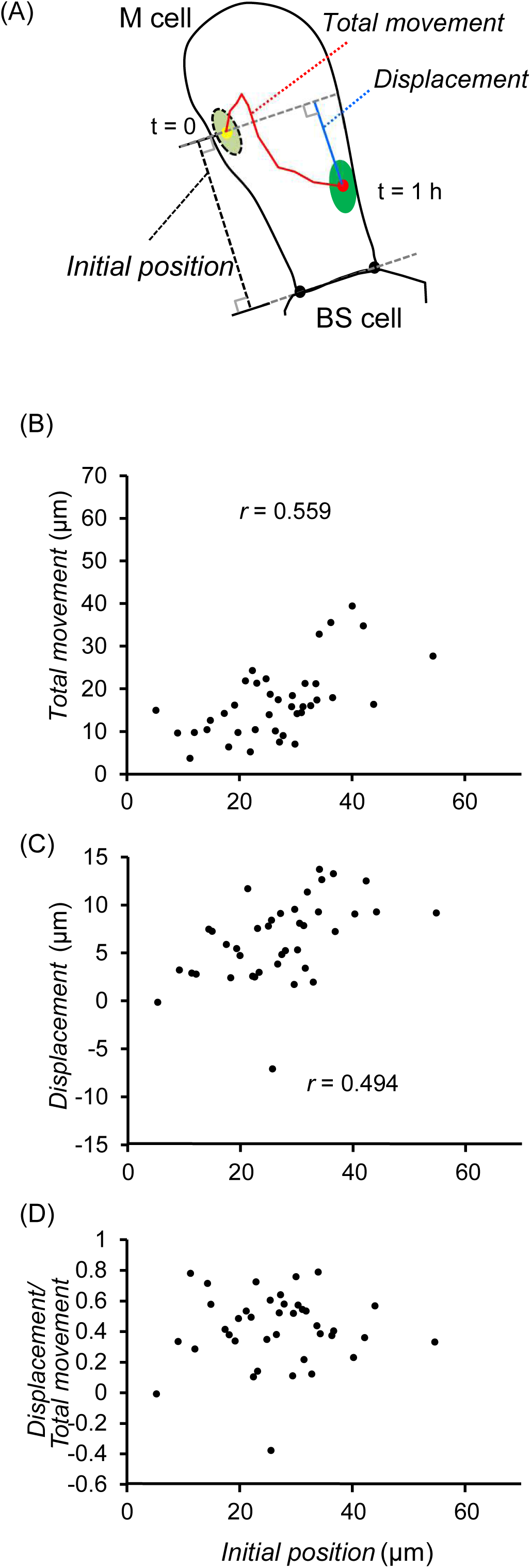
Correlation between the distance of M chloroplast from BS cell and motility of abaxial M chloroplast. (A) Schematic diagram showing M chloroplast positions before and after blue light irradiation and its distances traveled during the time. The total distance a chloroplast moved during the first hour of blue light irradiation is “*Total movement,*” and the linear distance to the BS cell side that the chloroplast moved for 1 h is “*Displacement*.” The linear distance between the chloroplast and the BS cell at the initial blue light irradiation is defined as the “*Initial position.*” (B) Correlations between *Initial position* and *Total movement*. (C) Correlations between *Initial Position* and *Displacement*. Negative *Displacement* values indicate that the chloroplast moved in the opposite direction from the BS cell side during the 1 h irradiation period. (D) Correlations between *Initial position* and *Displacement*/*Total movement*. *Displacement*/*Total movement* indicates migration efficiency to BS cells; the closer to +1, the more chloroplast movement is directed toward the BS cell side. The analysis was based on the data on chloroplast behavior shown in Fig. 2C (chloroplast number = 39). Pearson’s correlation test was conducted to evaluate the relationship, with the null hypothesis assuming no correlation between the two values (*p* < 0.05). The correlation coefficient was shown as *r* in each graph when a significant correlation was found.

### Necessity of BS cells for M chloroplast aggregative movements

We investigated whether BS cells were necessary for the movement of M chloroplasts toward BS cells by deactivating BS cell function. First, we attempted to selectively disrupt BS cells using a laser microbeam dissection system (Supplementary Figure S6). Because M and BS cells were adjacent, however, adjusting the light intensity to burst only BS cells was impossible. Therefore, as an alternative method for examining the behavior of M chloroplasts in the absence of BS cells, we focused on the differences in cell shape and attempted to remove only the BS cells. A ring structure of M and BS cells around the vein was observed in transverse sections (Figs. 4A, B). In contrast, M cells, BS cells, and vascular bundles were observed as stacked layers in longitudinal sections (Figs. 4A, C). The width of BS cells was larger than that of M cells in the transverse section (Fig. 4B). Therefore, a longitudinal section without BS cell contents was obtained by longitudinally cutting thinner than the BS cell and thicker than the M cell. Sections with BS cell contents were obtained by adjusting the thickness to 60 µm, while those without BS cell contents were obtained by setting the thickness to 40 µm. (Figs. 4D, E). In the M cells adjacent to the intact BS cells that maintained their cellular contents, the chloroplasts moved toward the BS cells and densely congregated at the junction between the M and BS cells under blue light irradiation from the adaxial side (Fig. 5A; Supplementary Movie S6). However, chloroplasts did not show typical aggregative movement in M cells adjacent to the dead BS cells without their cellular contents. Instead, they were aligned parallel to the direction of blue light incidence, moving away from the incident direction, which was reminiscent of the light avoidance response (Figs. 5B; Supplementary Movie S7). In sections with and without BS cell contents, when blue light irradiation was stopped, the M chloroplasts gradually dispersed to a state similar to that before blue irradiation. The finding that chloroplasts in M cells showed aggregative movement in response to blue light only when intact BS cells were adjacent to the M cells was also statistically confirmed (Fig. 5C). Although there was no significant difference in the speed of M chloroplast movement between with and without BS cell contents, M chloroplasts with BS cell contents significantly more oriented to the BS cells than without BS cell contents (Table 2). These results indicate that BS cells are necessary for the movement of M chloroplasts toward BS cells in response to blue light.

**Fig. 4.**
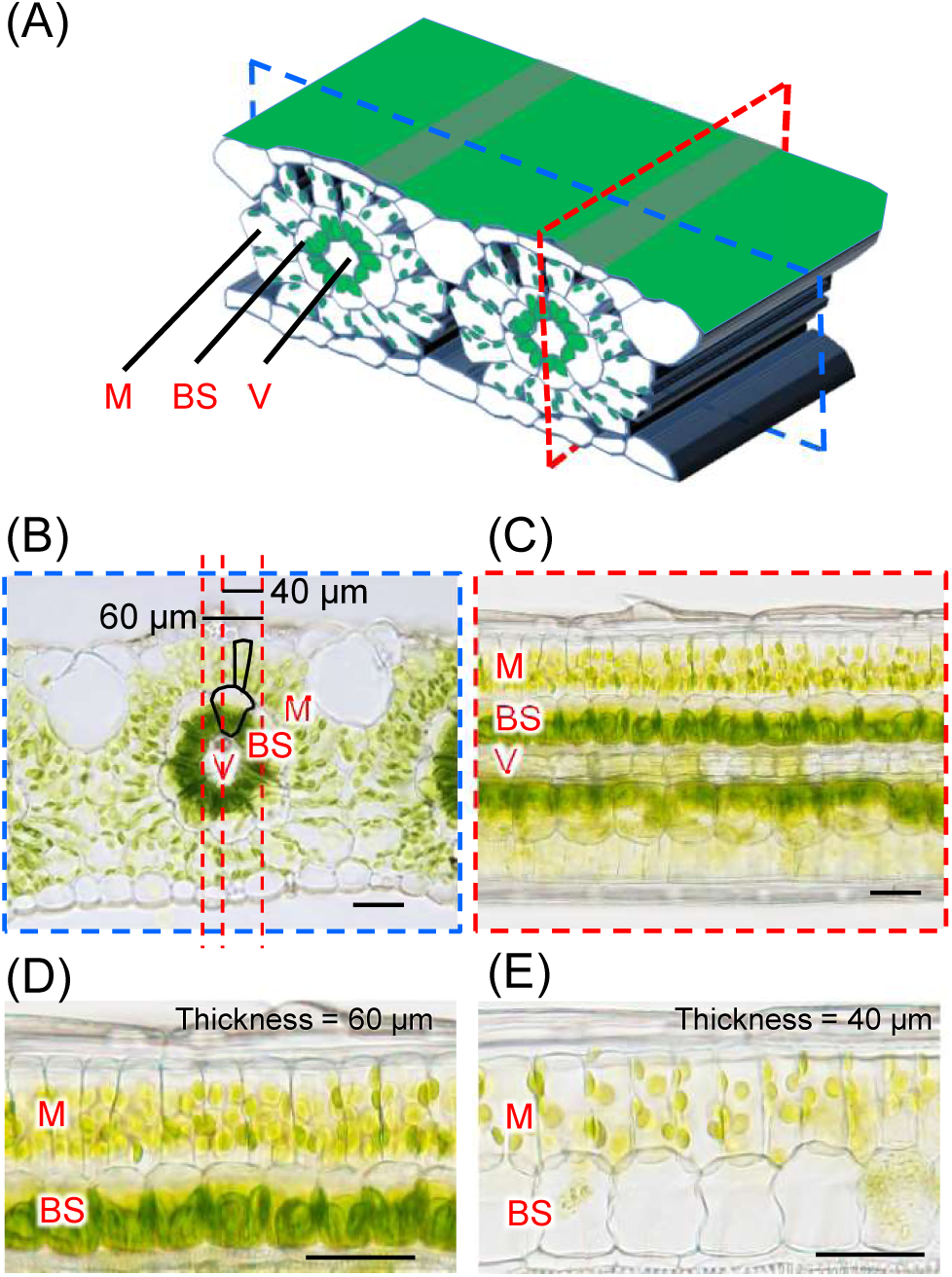
Sectioning of leaf tissue and removal of BS cell contents. (A) Schematic diagram of the leaf blade of finger millet. A transverse section was made by cutting the leaf blade perpendicular to the vascular bundle (blue dash line), and a longitudinal section was cut parallel to the vascular bundle (red dash line). Microscope images of a transverse section (B) and a longitudinal section (C). Since BS cells are larger than M cells, it is possible to cut only BS cells while leaving M cells intact, as shown by the red dashed line in the microscopic image (B). (D, E) adaxil side of longitudinal sections. (D) The longitudinal section with intact BS cells was obtained by sectioning at 60 µm thickness. (E) The longitudinal section without BS cell contents was obtained by sectioning at 40 µm thickness. M, mesophyll cell; BS, bundle sheath cell; V, vascular bundle. Bars in (B–E) = 50 µm.

**Fig. 5.**
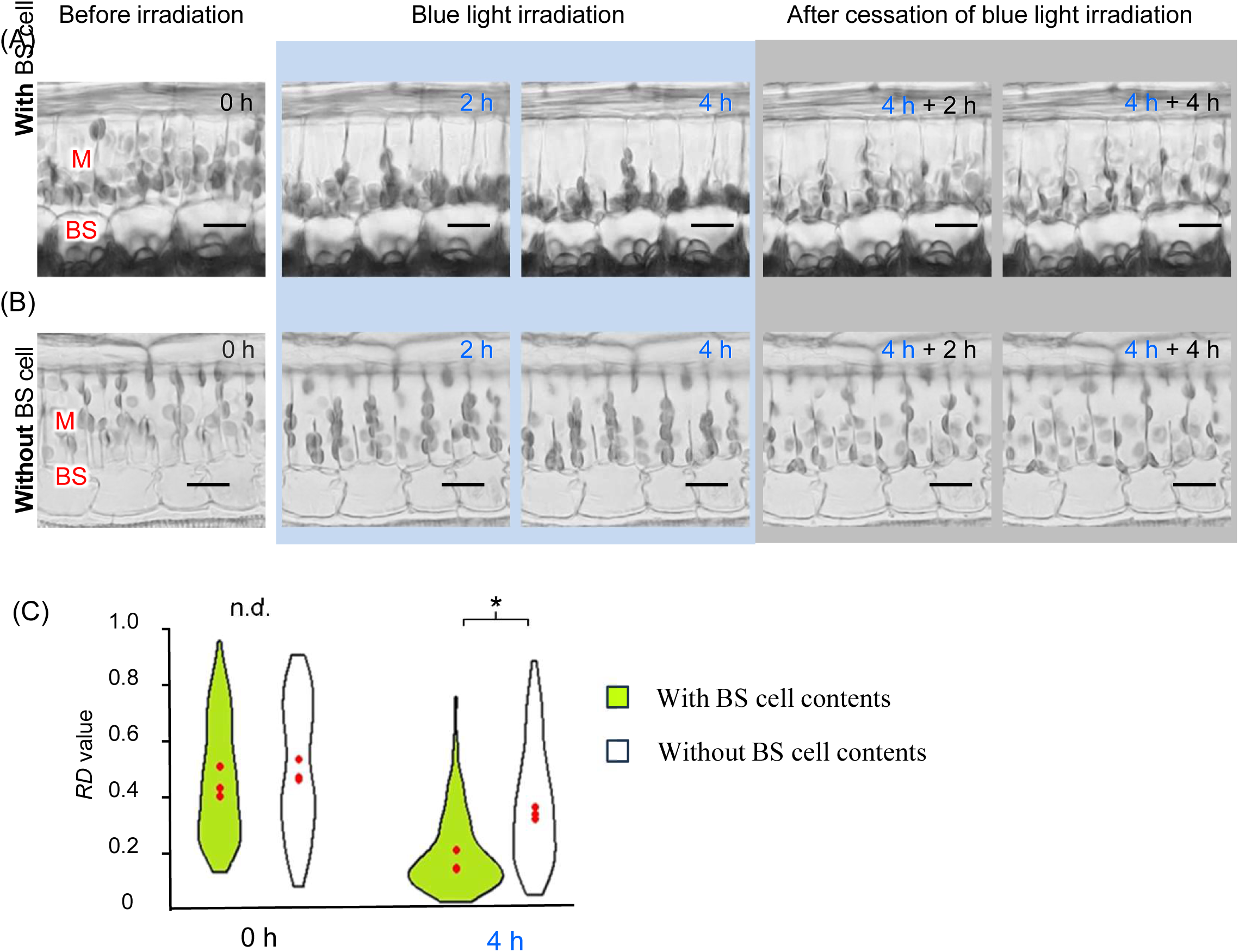
Comparison of adaxial M chloroplast movements with and without BS cell contents. Longitudinal sections with BS cell contents (A) or without the contents (B) were irradiated with blue light (500 µmol m^-2^ s^-1^) from the adaxial side. After 4 h of blue light irradiation, irradiation was stopped. The upper part of each image shows the adaxial side. Bars = 25 µm in (A, B). (C) Violin plots of M chloroplast relative distance (RD) from BS cells 0 and 4 h after blue light irradiation. Chloroplasts from three M cells per individual were measured, and three individuals were analyzed for each treatment (chloroplast number in each individual = 38–74). The width of each violin plot represents the probability density, and the red points in the violin plots indicate the *RD* median of each individual. Asterisks (*) indicate significant differences in the mean *RD* medians around each vascular bundle with and without BS cells (*p* < 0.05, Student’s *t*-test, *n* = 3).

**Table 2.** M chloroplast motility for 1 h blue light irradiation with or without adjacent BS cell contents.

|  | With BS<br>cell contents | Without BS<br>cell contents |  |
| --- | --- | --- | --- |
| Speed of chloroplast movement ($\mu\text{m min}^{-1}$) | 0.37±0.12 | 0.34±0.10 | |
| *Displacement/Total movement* | 0.41±0.22 | 0.29±0.23 | * |

Each value represents the mean ( ± SD). Asterisks indicate significant differences at $p < 0.05$ (Mann–Whitney $U$ test, chloroplast number = 44).

## Discussion

### Live leaf-section imaging allows continuous observation of chloroplast movement in the C_4_ leaf

Previous studies on chloroplast aggregative movement in C_4_ plants have compared chloroplast arrangements in leaf sections after chemical fixation under different conditions (Yamada *et al*., 2009, Maai *et al*., 2011b; Kato *et al*., 2023). In this study, we successfully established, for the first time, a live leaf-section imaging method that enables continual observation of chloroplast aggregative movements in M cells located inside the leaf tissues of the C_4_ plant finger millet (Fig. 1). The degree of M chloroplast arrangement toward the BS cells in response to intense blue light irradiation reached a maximum at approximately 4 h and was maintained until at least 16 h later (Supplementary Figures S3A; S4; Supplementary Movie S1). In previous experiments in which leaf fragments and fresh leaves were exposed to blue light, pronounced chloroplast aggregative movement occurred within 4 to 8 h, comparable to the movement observed in live leaf sections in the present study (Maai *et al*., 2011b; Kato *et al*., 2023). Furthermore, M chloroplasts returned to a dispersed arrangement after stopping blue light irradiation (Supplementary Figure S5; Supplementary Movie S3), indicating that the reversible responsiveness of chloroplast movement was retained in the leaf sections. Although our method requires careful consideration of the fact that samples were exposed to conditions different from the typical growth environment, such as loss of connectivity with surrounding cells and placement in a liquid-phase environment, it can be considered that the physiological response mechanisms related to chloroplast movement were maintained. Therefore, our leaf-section imaging technique, which allows for observing chloroplast movement inside the leaf tissue of C_4_ plants, is considered an effective method for analyzing chloroplast movement.

### M chloroplast aggregative movement occurs through communication with BS cells

M chloroplast aggregative movement is considered a physiological response associated with C_4_ photosynthesis, which functions between M and BS cells because the directionality of movement is toward the BS cells (Yamada *et al*., 2009; Taniguchi and Cousins, 2018; Kato *et al*., 2023). This movement has also been observed in various C_4_ plants (Kato *et al*., 2022). However, the relationship between M chloroplast aggregative movement and adjacent BS cells has not yet been confirmed. The analysis tracking individual M chloroplast movements toward adjacent BS cells demonstrated that the initiation time of M chloroplast movement induced by blue light would not be related to the distance from the BS cells at their initial position (Fig. 2B). Although M chloroplasts initially located farther from the BS cells tended to move more toward the BS cells (Fig. 3B-C), the efficiency of M chloroplast movement toward BS cells showed no correlation with the initial distance from the BS cells (Fig. 3D). These results did not clarify whether BS cells contributed to M chloroplast movement. To investigate the contribution of BS cells to the aggregation of chloroplasts in M cells, we selectively removed the BS cell content by adjusting the cutting angle and thickness (Fig. 4). The blue light-induced chloroplast aggregative movement was less likely to occur in M cells in the absence of adjacent intact BS cells; rather, chloroplasts showed a light avoidance response and were arranged parallel to the incident light direction (Fig. 5). This finding indicates that M chloroplast aggregative movement induced by blue light occurs through communication with BS cells, unlike the generally known in chloroplast photo relocation movement, which is a cell-autonomous response (Kagawa and Wada, 1994, 1996). Further research is needed to identify the molecular nature of BS cell signals that affect M cells’ chloroplast movement. Our leaf-section imaging technique, compatible with fluorescent molecular sensors, microbeam irradiation systems, and pulse amplitude-modulated fluorometry (PAM), will become an indispensable tool for investigating how chloroplast aggregative movement is involved in C_4_ photosynthesis.

## Supplementary data

Supplementary Movie S1. Live leaf-section imaging under continuous blue light irradiation from the adaxial side of the leaf section

Supplementary Movie S2. Live leaf-section imaging under no blue light irradiation

Supplementary Movie S3. Live leaf-section imaging under blue-light irradiation and after cessation of irradiation

Supplementary Movie S4. Live leaf-section imaging under continuous blue light irradiation from the abaxial side of the leaf section

Supplementary Movie S5. Tracking of M chloroplast movement

Supplementary Movie S6. Live leaf-section imaging of M chloroplasts with adjacent BS cell content under blue light irradiation from the adaxial side of the longitundinal leaf section

Supplementary Movie S7. Live leaf-section imaging of M chloroplasts without adjacent BS cells under blue light irradiation from the adaxial side of the longitundinal leaf section

## Acknowledgments

The author (Y.K.) would like to take this opportunity to thank the “Interdisciplinary Frontier Next-Generation Researcher Program of the Tokai Higher Education and Research System.”

## Author contributions

**Y.K. M.T. Y.S.** designed the study. Y.K. performed all the experiments, analyzed the data, and wrote the manuscript. M.T., T.O., and Y.S. supported the writing of the manuscript.

## Data Availability Statement

**For data available within the article or its supplementary materials:** All data supporting the findings of this study are available within the paper and supplementary materials published online.

## Abbreviations

BS: bundle sheath
M: mesophyll
RD: relative distance

