## Supplemental Figures for "Bundle sheath cell-dependent chloroplast movement in mesophyll cells of C_4_ plants analyzed using live leaf-section imaging"

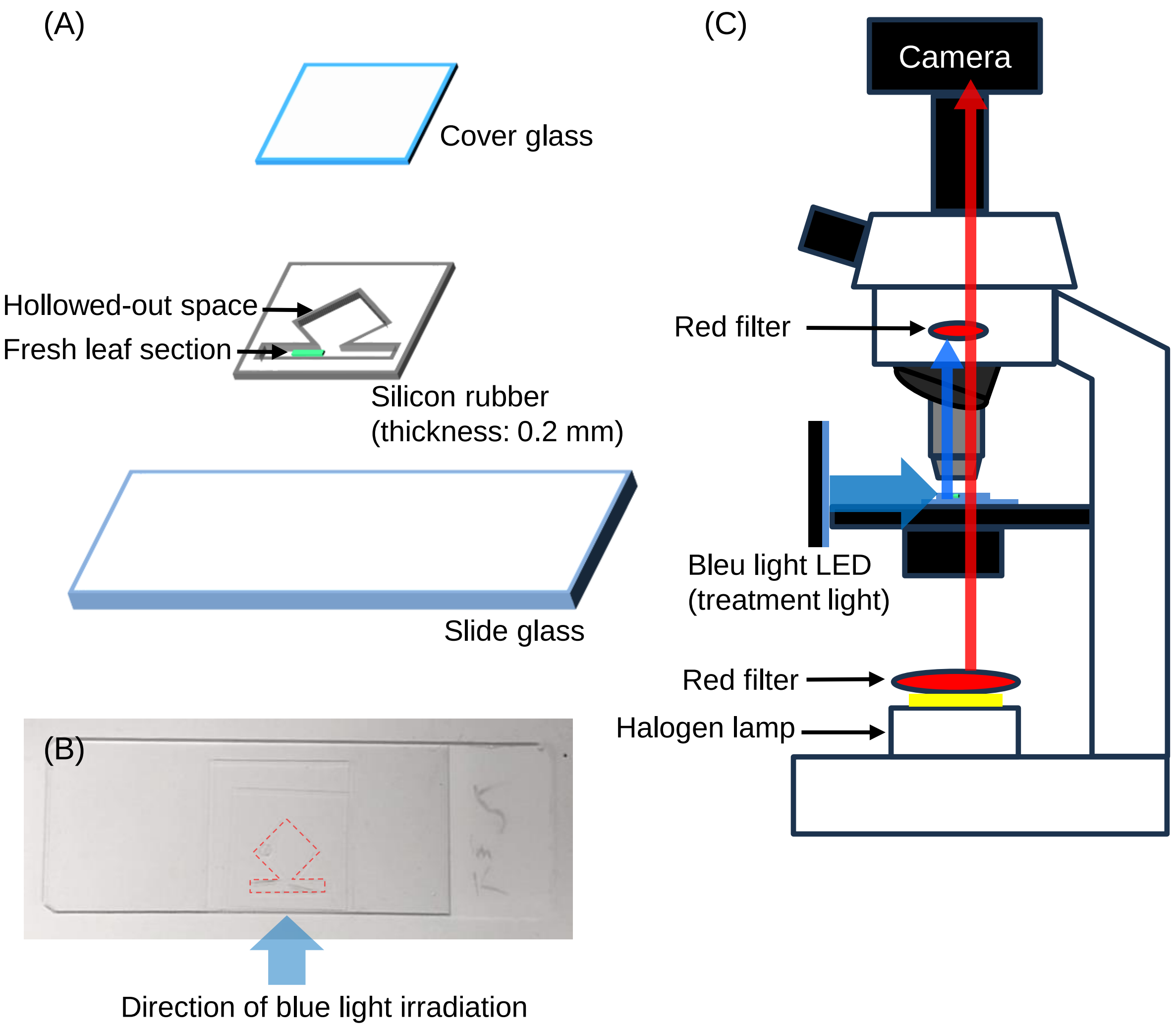


**Supplementary Figure S1. Schematic diagram of the live leaf-section imaging**

(A) Composition of the live leaf-section mount. (B) Top view of the mount and the direction of incidence of blue light. A transparent silicon rubber sheet with hollowed-out space for the groove was adhered to a slide glass. The leaf section was placed in the groove and enclosed with the cover glass. (C) Schematic diagram of the optical path. Red filters were set after the halogen lamp and in front of the camera in the optical path of the microscope light, respectively. Weak red light was used for observation (less than 1 μmol $m^{-2}$ $s^{-1}$).

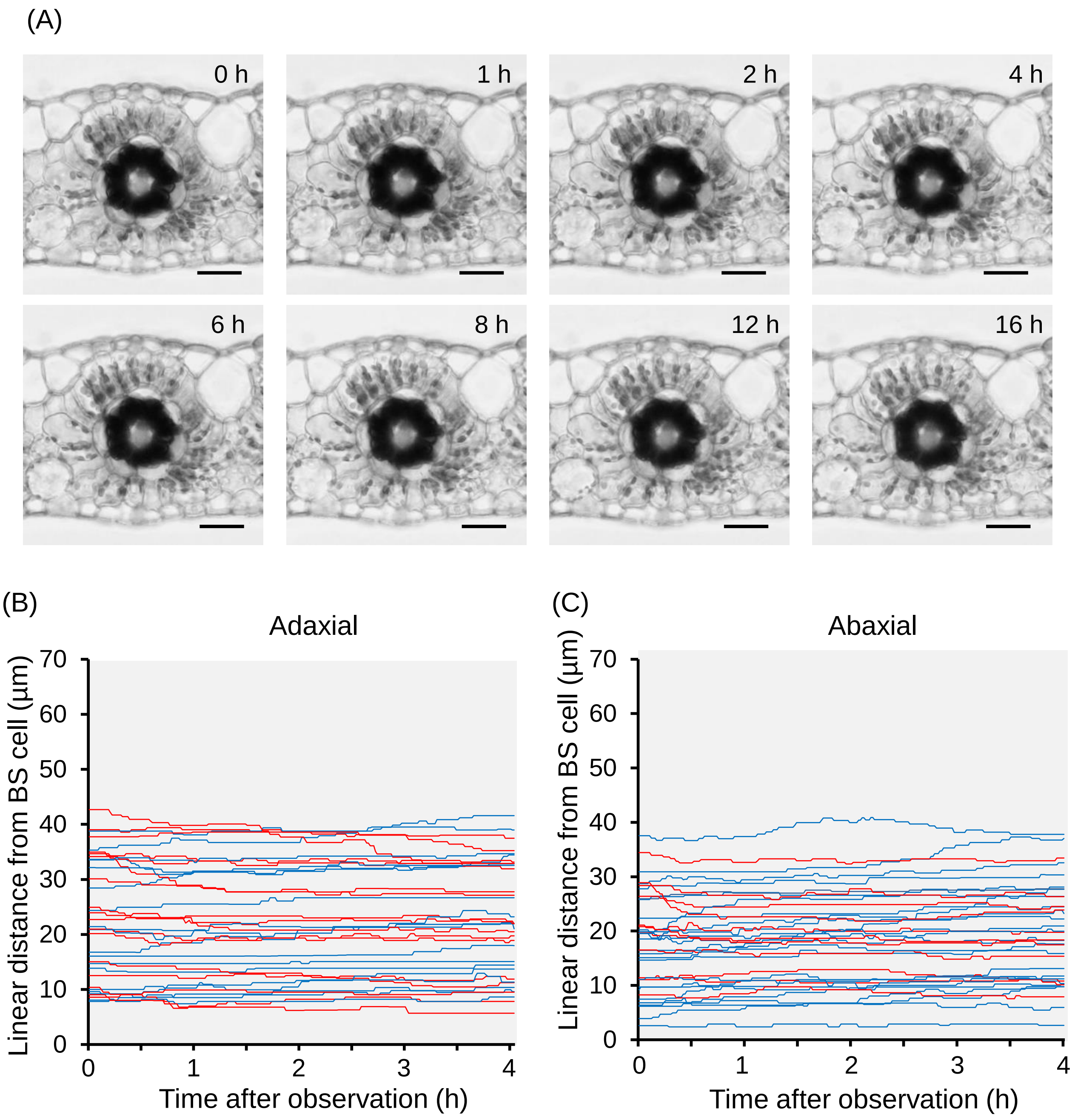


**Supplementary Figure S2. Temporal changes in the intracellular arrangement of chloroplasts in the absence of blue light**

(A) Time course images of transverse leaf tissue from the finger millet leaf blade. The transverse leaf section was not irradiated with blue light. The upper side of each image is the adaxial side. Bars = 50 µm. (B, C) Temporal changes in linear distance from BS cell in each M chloroplast were plotted every minute for 4 h without blue light irradiation. If chloroplasts could not be tracked due to overlapping each other or becoming out of focus, tracking was terminated at that point. Red lines represent chloroplasts that moved closer to BS cells, and blue lines represent those that moved farther from BS cells after 4 h of blue light irradiation compared to their initial position at the initial observation (t = 0). Their line colors were determined based on the end point of tracking for chloroplasts, which could not be tracked until 4 h of blue light irradiation. Over ten chloroplasts were randomly selected for monitoring from three individuals' adaxial and abaxial sides, respectively (chloroplast number in the adaxial side = 35, those in the abaxial side = 36).

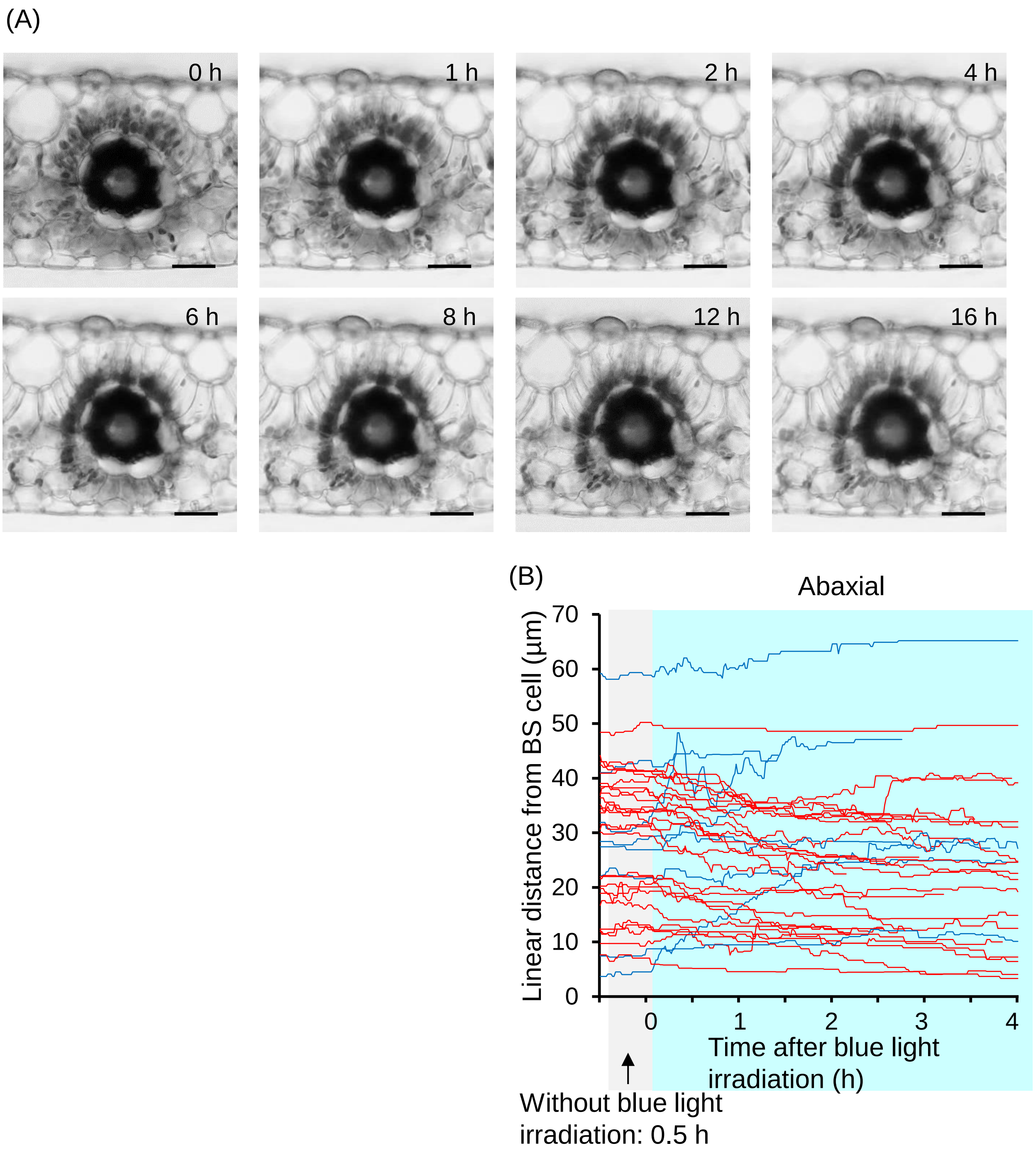


**Supplementary Figure S3. Temporal changes in the intracellular arrangement of chloroplasts under blue light irradiation**

(A) Time course images of transverse leaf tissue from the finger millet leaf blade. The leaf section was irradiated with blue light (500 µmol $m^{-2}$ $s^{-1}$) from the adaxial side. The upper side of each image is the adaxial side. Bars = 50 µm. (B) Temporal changes in linear distance from BS cell in each M chloroplast on the abaxial side were plotted every minute for 4 h under blue light irradiation. If chloroplasts could not be tracked due to overlapping each other or becoming out of focus, tracking was terminated at that point. Red lines represent chloroplasts that moved closer to BS cells, and blue lines represent those that moved farther from BS cells after 4 h of blue light irradiation compared to their initial position at the initial irradiation (t = 0). Their line colors were determined based on the end point of tracking for chloroplasts that could not be tracked until 4 h blue light irradiation. Over ten chloroplasts were randomly selected for tracking from the abaxial side of leaf sections in three individuals (chloroplast number = 35). See also Figure 2B showing the adaxial side changes.

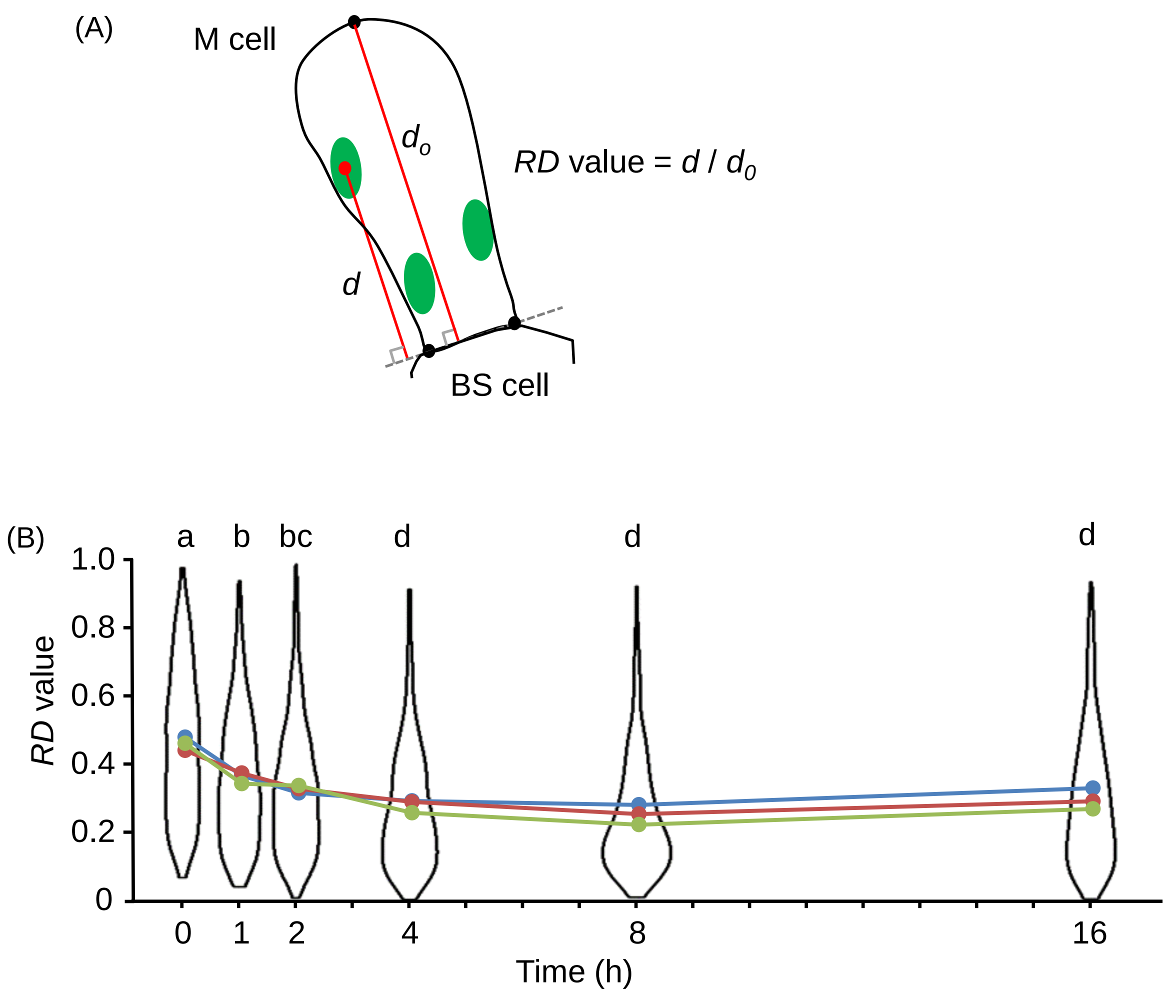


**Supplementary Figure S4. Temporal changes in the degree of chloroplast aggregative arrangements under blue light irradiation**

(A) Schematic diagram of relative distance (*RD*) value. (B) Violin plots representing the *RD* of M chloroplasts. Three groups of M cells surrounding a vascular bundle were analyzed, and each cell group was derived from a different plant individual (chloroplast numbers in each individual = 212—313). The width of each violin represents the probability density. The same color indicates the median value of the same individual at each time. Different letters above the violin plots indicate a significant difference in the mean of the *RD* medians around each vascular bundle at $p < 0.05$ (Tukey–Kramer method, $n = 3$).

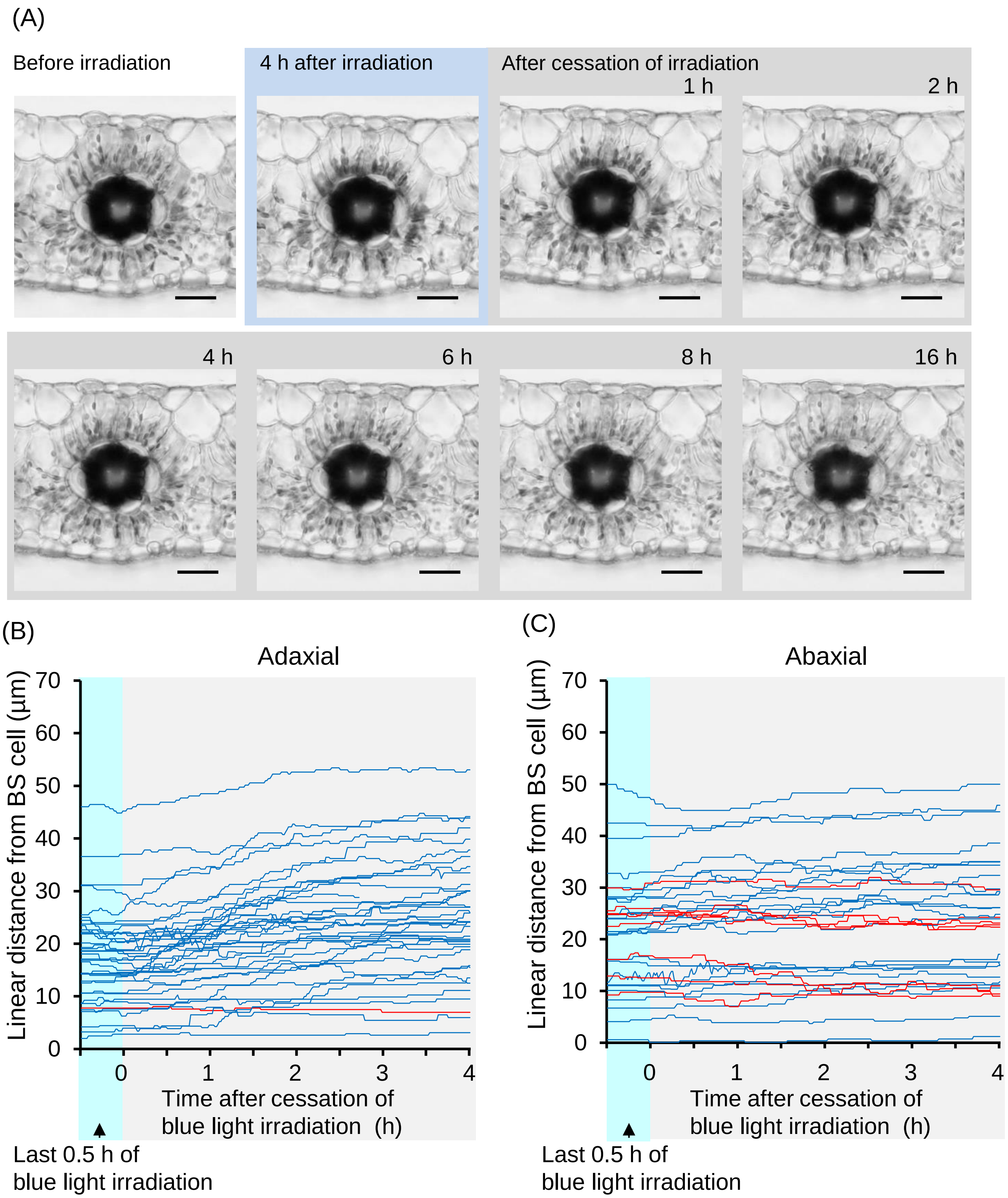


**Supplementary Figure S5. Temporal changes in the intracellular arrangement of chloroplasts under blue light irradiation and after cessation of the irradiation**

(A) Time course images of transverse leaf tissue from the finger millet leaf blade. The transverse leaf section was irradiated with blue light (500 μmol $m^{-2}$ $s^{-1}$) from the adaxial side. After 4 h of the blue light irradiation, irradiation was stopped. The upper side of each image is the adaxial side. Bars = 50 μm. (B, C) Temporal changes in linear distance from BS cells in each chloroplast were plotted every minute from the last 0.5 h of 4 h of blue light irradiation to 4 h after the cessation of the irradiation. If chloroplasts could not be tracked due to overlapping each other or becoming out of focus, tracking was terminated at that point. Red lines represent chloroplasts that moved closer to BS cells, and blue lines represent those that moved farther from BS cells after 4 h compared to their initial position at the cessation of irradiation (t = 0). Their line colors were determined based on the end point of tracking for chloroplasts, which could not be tracked until 4 h of irradiating blue light. Over ten chloroplasts were randomly selected for monitoring from three individuals' adaxial and abaxial sides, respectively (chloroplast number in the adaxial side = 39, those in the abaxial side = 33).

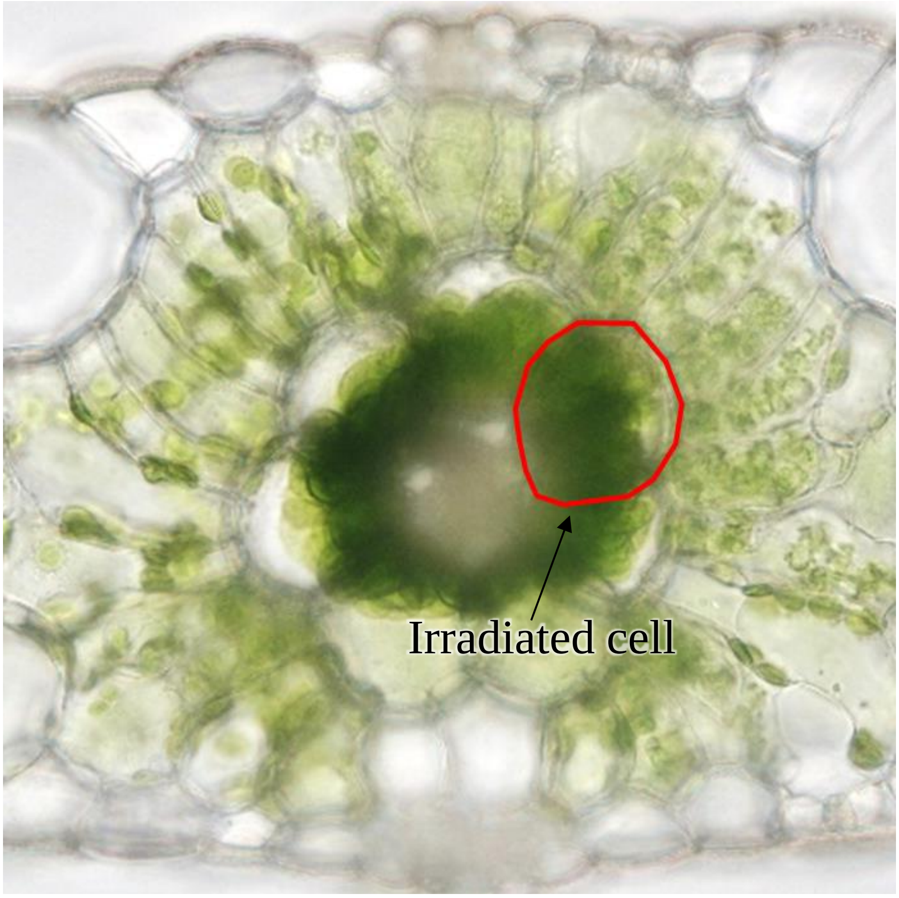


**Supplementary Figure S6. Disruption of BS cell by laser microbeam irradiation**

Images of leaf tissue transverse sections irradiated with a laser microbeam using a PALM Combi system (Zeiss, Germany). The center of the BS cell was irradiated with a microbeam to disrupt the cell selectively. However, the chloroplasts in the M cells neighboring the BS cells irradiated with the microbeam ruptured and no longer moved.
